# Attention hinges on tuned normalization strength within human visual cortex

**DOI:** 10.1101/515254

**Authors:** Ilona M. Bloem, Sam Ling

## Abstract

Although attention is known to increase the gain of visuocortical responses, its underlying neural computations remain unclear. Here, we used fMRI to test the hypothesis that a neural population’s ability to be modulated by attention is dependent on divisive normalization. To do so, we leveraged the feature-tuned properties of normalization and found that visuocortical responses to stimuli sharing features normalized each other more strongly. Comparing these normalization measures to measures of attentional modulation, we discovered that subpopulations that exhibited stronger normalization also exhibited larger attentional benefits. In a converging experiment, we demonstrated that attentional benefits were greatest when a subpopulation was forced into a state of stronger normalization. We propose a tuned normalization model of attention that parsimoniously accounts for many properties of our results, suggesting that the degree to which a subpopulation exhibits normalization plays a role in dictating its potential for attentional benefits.

## Introduction

Neural processing is surprisingly efficient. Although our environment is brimming with information, our cognitive system adeptly regulates competition between neural representations– all competing for visual awareness. A growing body of evidence suggests that this is made possible, in part, by recruiting a seemingly ubiquitous neural computation, known as divisive normalization, which can regulate the relative strength between competing representations^1–4^. Under normalization, the response to a stimulus is modulated by the summed activity generated by the stimulus itself, along with pooled neighboring responses. This computation crucially supports a number of functions, including regulating the dynamic range of neural responses^2,3,5,6^. Models of divisive normalization have long served as cornerstone principles for computational accounts of early vision, and generalize to a variety of other sensory modalities^3,7,8^ and cognitive processes^5,9–14^. Interestingly, normalization is also believed to be modulated by contextual influences, whereby visual features that are similar tend to normalize each other more than those that are dissimilar^10,15–18^. This feature-tuned aspect has been theorized to play an active role in reducing redundant sensory information^17,19–22^. Our visual environment is comprised of statistical biases between image properties, whereby neighboring features belonging to the same object are most likely to be similar. Feature-tuned normalization incorporates these inherent dependencies, acting as a form of neural information compression by reducing statistical biases in natural images, thereby deprioritizing the processing of redundant representations^17,21,22^.

While tuned normalization may play a role in the bottom-up enhancement of potentially relevant information in a visual scene, we ultimately rely on top-down attentional systems to selectively enhance a small subset of that information for prioritized processing, from moment to moment. One of the most well-documented ways that attention enhances relevant information is by means of an increase in the gain, or ‘strength’, of the behavioral^23–26^ or neural response^5,27–30^. Interestingly, prominent computational models have theorized that normalization and attention are tightly linked, whereby attentional modulation within visual cortices is dependent on divisive normalization^5,31,32^. These models propose that attention can alter the balance between the stimulus activity and the summed activity of the normalization pool, loosening the current state of gain control, and thereby resulting in an increased neural response. Normalization accounts of attention have traditionally hinged on three key components: the locus of attention, the size of the stimulus, and the size of the attentional window^5,28,33^. However, these models consider normalization to be feature-agnostic, allowing an equal contribution of all information, regardless of feature similarities. The notion that attention modulation could additionally depend on a fourth component, incorporating the feature-selective nature of normalization has some support in animal studies, with single-unit recordings in macaques suggesting that the contribution of tuned normalization can explain attention biases of competition between multiple stimuli within a receptive field^10^.

In this study, we used functional magnetic resonance imaging (fMRI) to test the hypothesis that attention-driven modulation of the gain of responses within human visual cortex is made possible, in part, by a release from feature-tuned normalization. We approach the problem by first devising an efficacious, voxel-by-voxel population measure of the feature-tuned aspects of normalization within early visual cortex, during passive viewing. To do so, we exploited a phenomenon known as sub-additivity, a signature property of normalization wherein the population responses to images comprised of two superimposed stimuli tend to fall short of the linear sum of the response to each stimulus independently^34–39^. We discovered potent tuned normalization within human visual cortex: superimposed stimuli sharing the same features were more strongly normalized than stimuli that differed in their features. Armed with a population measure of feature-tuned normalization, we set out to test the hypothesis that attentional modulation is partially driven by tuned normalization. If normalization truly governs attentional modulation, we reasoned that attention-driven gain changes would be greater when a neural subpopulation within early visual areas exhibits stronger normalization. Indeed, in our second experiment we reveal that tuned normalization is tightly linked to an independent measure of attentional modulation. Leveraging population-wide heterogeneities in BOLD responses for both normalization and attention measures, we found that subpopulations that exhibited stronger normalization also exhibited larger attentional benefits. In a third converging experiment, we directly manipulated spatial attention, while simultaneously measuring population activity under different states of normalization. In doing so, we found that attentional benefits are greater when the population is put under stronger normalization. Finally, we introduce a variant of the normalization model of attention, which reveals that the incorporation of feature-tuned normalization nicely captures our results –a neural population’s capability for attentional benefits appears contingent upon normalization, whereby the degree to which a population can normalize itself results in greater potential for release from normalization driven by attention.

## Results

### Sub-additivity as a signature of tuned normalization

We first set out to obtain a population measure of visuocortical responses under different states of normalization. Specifically, in addition to a well-known untuned, feature agnostic component, does sub-additivity show a signature of a tuned, or feature-selective component? We leveraged the fact that population responses to images comprised of superimposed visual stimuli are not simply the linear sum of the response to each stimulus independently^34–37^; instead the response typically exhibits a property known as sub-additivity–a phenomenon nicely captured by contrast normalization. This is believed to emerge due to the compressive nature of normalization, which acts to nonlinearly limit the overall response to the stimuli. To assess the influence of tuned and untuned normalization on population responses within human visual cortex, we leveraged an fMRI noise-masking technique^24,35,37^, which allowed us to test the degree to which BOLD responses within early visual cortex exhibit sub-additivity, depending on stimulus feature similarity. To tap into the sub-additive nature of tuned normalization, we constructed stimuli that were composed of linearly summed pairs of oriented bandpass-filtered noise gratings (outer diameter 15°; inner diameter 3°; at 50% Michelson contrast; spatial frequencies between 2-3 cycles/°; orientation bandwidth of 10°). Importantly, images were constructed using pairs of stimuli combined in either an *orthogonal* (different features) or a *collinear* configuration (similar features; **Figure 1a**). We measured BOLD responses to these collinear and orthogonal stimuli configurations in separate blocks during an fMRI session, while participants performed a demanding task at fixation, finding a target in a rapid letter stream, and ignoring the stimuli presented in the periphery (**Figure S1**). Additionally, in a separate set of scans we measured the BOLD response to each individual component that comprised the overlaid stimuli, and doubled this obtained response in order to create a hypothetical additive sum. The sub-additive deviation from the hypothetical sum for both the collinear and orthogonal configurations served as our measure of untuned normalization, while comparing the difference between the configurations served as our measure for an additional tuned component.

**Figure 1.**
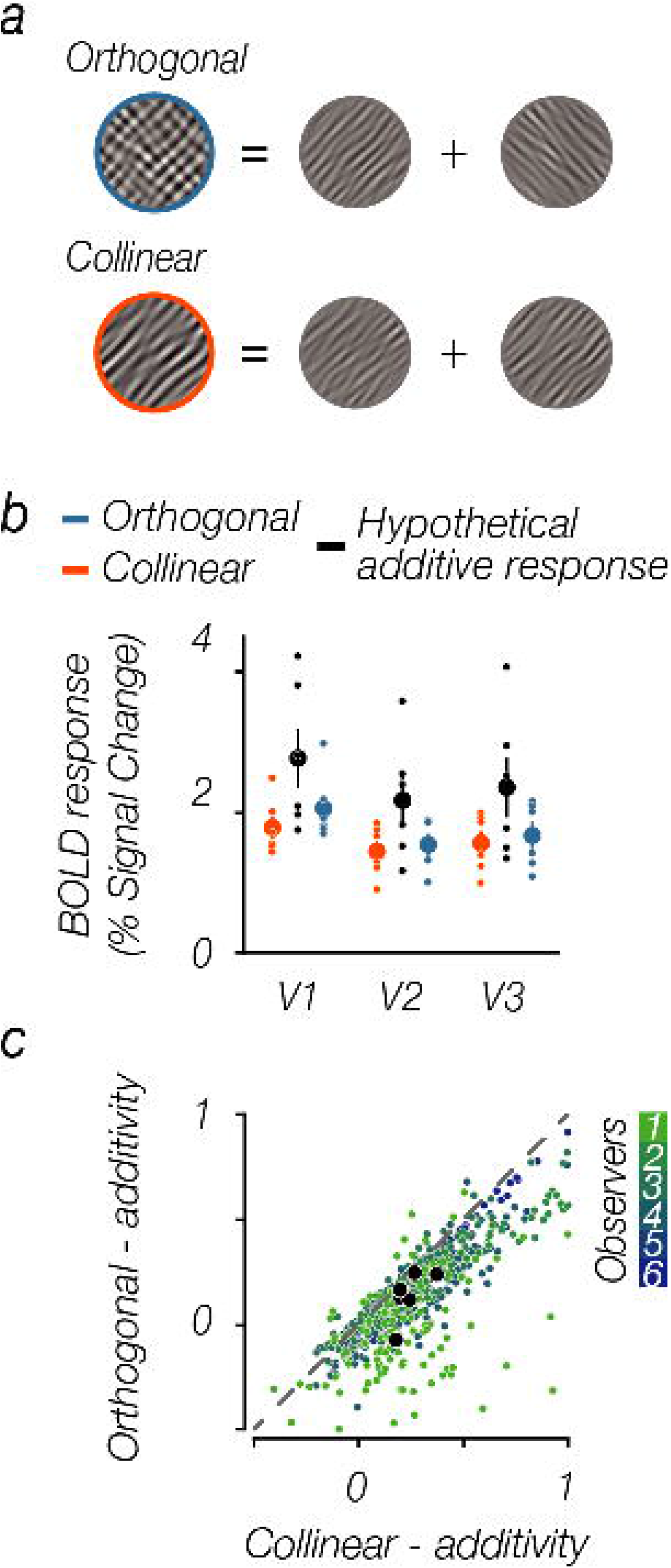
Sub-additivity as a measure of tuned normalization. a. Schematic of stimuli used to measure sub-additivity. Two oriented stimuli components (outer diameter 15°; inner diameter 3°, at 50% Michelson contrast, spatial frequencies between 2-3 cycles/°; orientation bandwidth of 10°) were linearly summed in either an orthogonal (top; blue) or collinear (bottom; orange) configuration, resulting in a full contrast stimulus. Stimuli modified for illustrative purposes. b. Average BOLD responses across observers for orthogonal (blue) and collinear (orange) configurations. A hypothetical additive response (black) was created by doubling the BOLD response evoked by an individual stimulus component. Dots represent individual observers; error bars denote ±1S.E.M.. c. Voxel-wise relationship between sub-additivity measures for both configurations in V1. BOLD responses were normalized for each participant. Small colored dots indicate individual voxels; larger black dots represent the whole ROI average per observer.

Orthogonal and collinear stimuli configurations both exhibited sub-additivity across early visual areas (V1-V3), with the measured BOLD response of both stimulus configurations being lower than the hypothetical additive sum of the individually measured components (paired one sided t-test; orthogonal sub-additivity in V1: *t(5)* = 2.07, *p* = 0.0465, V2: *t(5)* = 2.70, *p* = 0.0215, and V3: *t(5)* = 2.46, *p* = 0.0287; collinear sub-additivity in V1: *t(5)* = 2.89, *p* = 0.0171, V2: *t(5)* = 2.87, *p* = 0.0175, and V3: *t(5)* = 2.58, *p* = 0.0247; **Figure 1b & S1**). Furthermore, the responses demonstrated robust feature-tuned normalization as well, whereby stimuli comprised of collinear orientations were more sub-additive, and thus more strongly normalized, than stimuli that contained orthogonal orientations (paired two sided t-test; V1: *t(5)* = 5.97, *p* = 0.0019, V2: *t(5)* = 3.82, *p* = 0.0123, and V3: *t(5)* = 3.44, *p* = 0.0184). While BOLD responses to either stimuli configuration across visual areas were fairly consistent in the degree to which they exhibited sub-additivity (untuned normalization), the magnitude of the feature-tuned aspect of normalization seemed to decrease in strength along the visual hierarchy. This orientation-tuned aspect of normalization was strongest within primary visual cortex –a region shown to be most precisely tuned to orientation content^40–42^, and became less apparent as we moved up the visual hierarchy, consistent with a shift in the preferred feature space. We next explored the degree of dependency, from voxel-to-voxel, of the deviation from additivity for both orthogonal and collinear stimuli configurations within V1. Although there is heterogeneity in the magnitude of sub-additivity between voxels within a region, comparing the magnitude of sub-additivity for collinear and orthogonal stimulus configurations revealed a consistent pattern, in that the collinear configuration evoked lower BOLD responses compared to the orthogonal configuration for almost all voxels within an area (**Figure 1c**).

Importantly, the differences in BOLD responses evoked by the two stimuli configurations were not driven by differences in the image statistics, nor could they be explained by basic first-order visual response properties. A Fourier analysis confirmed that while the orientation content between the stimuli configurations differed, the overall power was comparable (**Figure 2a**). Furthermore, a V1-based energy detection model^43–48^ that only incorporated untuned divisive normalization also fell short of accounting for these results. In this model, we estimated the amount of contrast energy each class of images evoked by applying a linear Gabor wavelet decomposition that described tuning along the dimensions of space, phase, orientation, and spatial frequency^43,44^ (**Figure 2b**). 500 unique images for both collinear and orthogonal stimuli configurations were passed through the model, which resulted in a measure of contrast energy for each image, contained in quadrature wavelet pairs. After combining all wavelets across space and spatial frequency scales, a measure of contrast energy evoked by each orientation channel remained. The model output undergoes divisive normalization, effectively acting as a contrast gain control operator^2,6,49^. Stimulus energy was demeaned and normalized to 1. A bootstrap analysis indicated that there is no difference between the two simulated stimulus energy distributions, indicating that these images have the same stimulus energy when applying untuned normalization (95% confidence interval = [-0.026, 0.019]; **Figure 2c**). Importantly, however, a difference in stimulus energy was observed when we built an orientation-tuned component into the normalization model (bootstrapped 95% confidence interval = [0.362; 0.4035]; **Figure 2d**), indicating that the observed differences in sub-additivity of the BOLD responses between the two stimuli configurations can be driven by tuned normalization.

**Figure 2.**
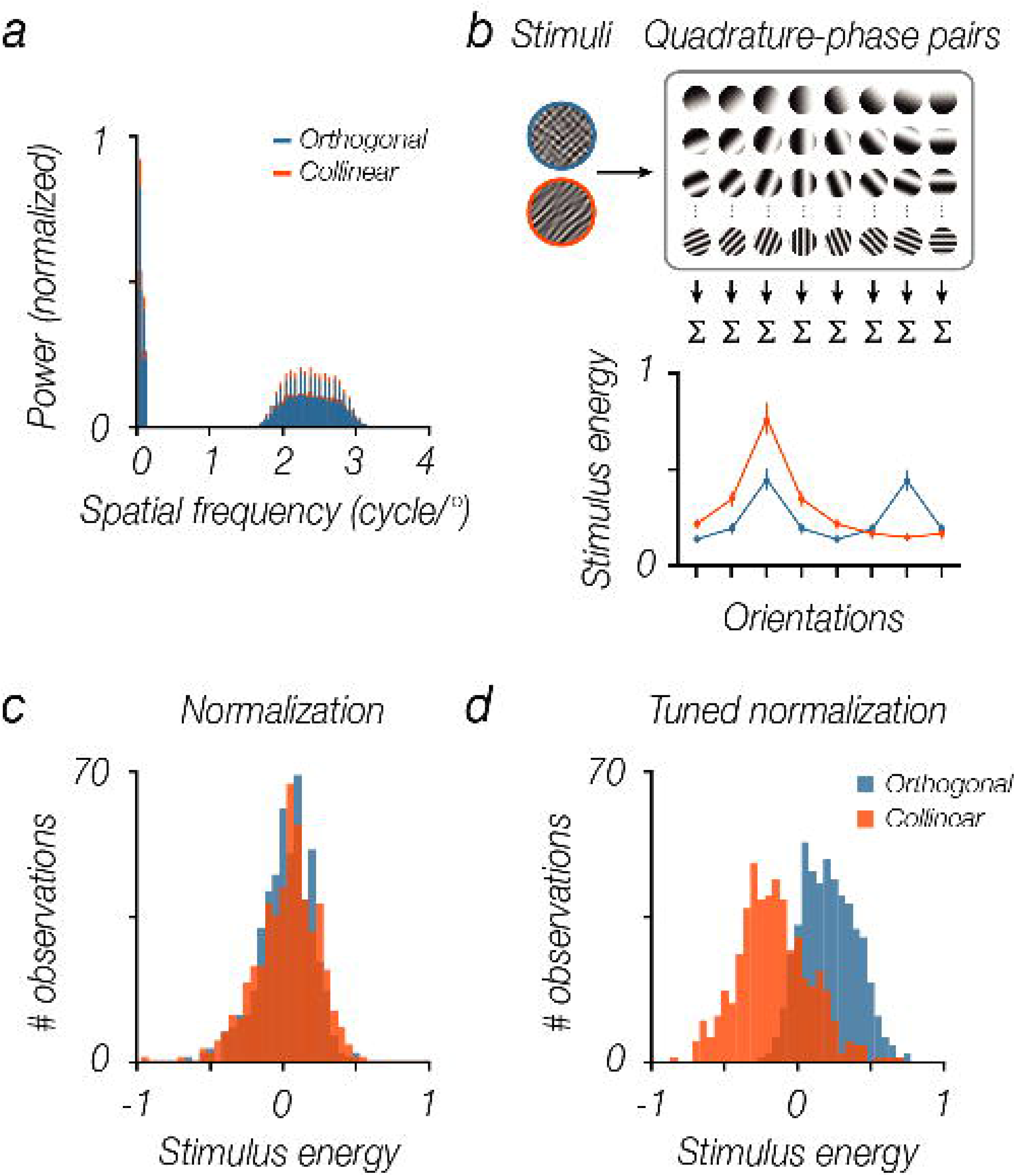
Image statistics between collinear and orthogonal stimuli did not differ, a. Average normalized power obtained with a 2D Fourier transform of 500 images within each stimulus configuration (collapsed across phase). Error bars denote ±1 S.E.M.. b. Energy detector model. Stimuli were convolved with a set of Gabor filters across space (centered on every pixel of the image), phase (2 phases to create a quadrature pair), spatial frequency (36 scales, ranged between 0.5-4 cycles/°), and orientations (8 orientation channels). The quadrature-phase pairs were squared, summed, square-rooted, and normalized to 1, to generate a complex cell response at the 8 orientation channels for each image. Error bars denote ± 1 S.E.M.. c. To account for contrast saturation, divisive normalization was applied to the total stimulus energy obtained from the filter outputs, and we combined all orientation channels to obtain one stimulus energy value per image. As is evident, the distributions of stimulus energy for both stimuli configurations are highly comparable when divisive normalization is applied. d. Incorporating a tuned component to the normalization term results in a separation of the energy distributions in which collinear stimuli have a relatively lower stimulus energy.

### Attentional modulation is related to tuned normalization strength

Leveraging our ability to measure feature-tuned normalization within human visual cortex, we then set out to test our main hypothesis: Does attention optimize information processing by modulating divisive normalization? Our previous experiment demonstrated that collinear stimuli configurations were more sub-additive as they evoked lower BOLD responses compared to orthogonal stimuli. Importantly, while responses to each stimulus configuration varied substantially, this difference between collinear and orthogonal configurations was consistent for almost all voxels within an area (**Figure 1c**). In a second experiment, we examined the degree of voxel-wise dependency between this measure of tuned normalization and an independent measure of attention modulation. If normalization and attentional modulation interact, we predict that those sub-populations that are more strongly normalized should also exhibit the highest potential for attentional modulation.

Tuned normalization strength was quantified as the difference between the mean BOLD responses to orthogonal and collinear stimuli blocks. Collinear stimuli evoked weaker BOLD responses compared to orthogonal stimuli, resulting in a positive difference for all regions of interest (**Figure 3a**; two-sided t-test; V1: *t(5)* = 5.97, *p* = 0.0019, V2: *t(5)* = 3.82, *p* = 0.0123, and V3: *t(5)* = 3.44, *p* = 0.0184). To measure attentional modulation, we assessed BOLD responses while participants viewed orientation bandpass–filtered noise gratings (outer diameter 15°); inner diameter 3°; at 50% Michelson contrast; spatial frequencies between 2-3 cycles/°; orientation bandwidth of 10°). While viewing these stimuli, participants were asked to either covertly attend towards the stimulus, performing a fine orientation discrimination task (*Attended condition*), or perform a demanding task at fixation, which drew attention away from the oriented stimulus (*Unattended condition*). Note that the visual stimulation was identical in both conditions, the only difference being the task observers performed (**Figure S2**). Attention modulation was defined as the difference between BOLD responses to attended and unattended stimuli blocks. Consistent with previous findings^24,50–52^, striate and extrastriate cortex exhibited robust attentional modulation (**Figure 3b**; two-sided t-test; V1: *t(5)* = 4.44, *p* = 0.0068, V2: *t(5)* = 3.85, *p* = 0.0120, and V3: *t(5)* = 4.07, *p* = 0.0096).

**Figure 3.**
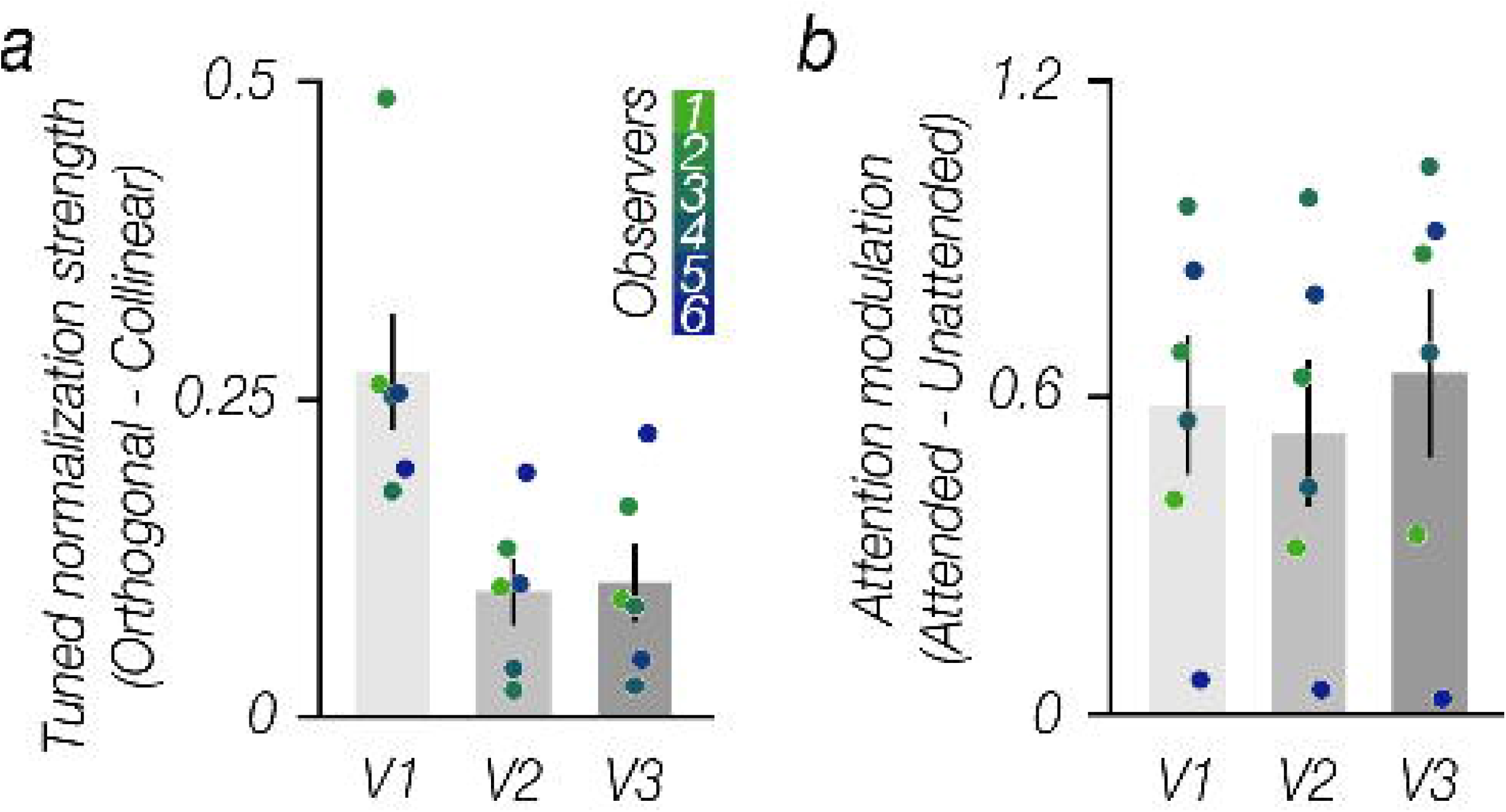
Measures of tuned normalization strength and attention modulation. a. Tuned normalization strength reflects the difference between BOLD responses evoked by orthogonal and collinear stimuli blocks (% signal change). The orthogonal configuration elicited larger BOLD responses compared to the collinear configuration across V1-V3. b. Attention modulation reflects the difference in BOLD response between attended and unattended stimuli blocks (% signal change). We observed large attention modulation across V1-V3. Error bars denote ± 1 S.E.M., colored dots indicate individual observers.

While these results reflect the average response across the entire region of interest (ROI), the magnitude of tuned normalization strength varied substantially from voxel-to-voxel within each visual area, suggesting heterogeneity across the population (**Figure 1c**). Leveraging this population-wide heterogeneity in neural responses in both our attentional modulation and tuned normalization measures, our results revealed that subpopulations that exhibit the strongest tuned normalization also possess the greatest attentional benefits, across visual areas (**Figure 4**; two-sided t-test of Fisher Z transformed Spearman correlations; V1: *t(5)* = 4.30, *p* = 0.008, V2: *t(5)* = 2.96, *p* = 0.0315, and V3: *t(5)* = 5.14, *p* = 0.004). To ensure that our results reflected a true relationship between normalization and attention, rather than being driven by differences in spurious factors, such as the signal-to-noise ratio (SNR), our analyses were restricted to the top 25% most visually responsive voxels within V1-V3, selected using an independent functional localizer. Note that the relationship persists when all voxels within a respective region are used in the analyses (V1: *t(5)* = 4.30, *p* = 0.008, V2: *t(5)* = 3.06, *p* = 0.0286, and V3: *t(5)* = 2.80, *p* = 0.04). In addition, we examined whether the relationship between tuned normalization strength and attentional modulation was still evident even when we broke down our stringent voxel selection into four bins, according to ranked goodness of fit (R^2^) of responses to the visual localizer (**Figure 4b**). While the observed correlation within V1 and V2 was not driven by differences in R^2^ of the localizer scans as a similar relationship persisted in each bin, within V3 the correlation does seem driven by voxels that had a higher R^2^ (two-sided t-test of Fisher Z transformed Spearman correlations; V1: Q1 *t(5)* = 4.01, *p* = 0.010, Q2 *t(5)* = 3.35, *p* = 0.020, Q3 *t(5)* = 3.66 *p* = 0.015, Q4 *t(5)* = 5.94, *p* = 0.002; V2: Q1 *t(5)* = 2.51, *p* = 0.054, Q2 *t(5)* = 2.35, *p* = 0.066, Q3 *t(5)* = 2.44 *p* = 0.059, Q4 *t(5)* = 2.12, *p* = 0.087; and V3: Q1 *t(5)* = 6.46, *p* = 0.001, Q2 *t(5)* = 4.49, *p* = 0.007, Q3 *t(5)* = −0.24 *p* = 0.817, Q4 *t(5)* = 1.98, *p* = 0.105). The less pronounced relationship in V3 is likely driven by the reduced heterogeneity and overall magnitude of tuned normalization strength observed in this visual area (**Figure 4a**). Taken together, leveraging the heterogeneity of population responses for attentional modulation and tuned normalization strength, our results reveal a tight link between these two measures, which was strongest in primary visual cortex, suggesting that a neural subpopulation’s potential to increase its attentional gain is dependent on its tuned normalization strength.

**Figure 4.**
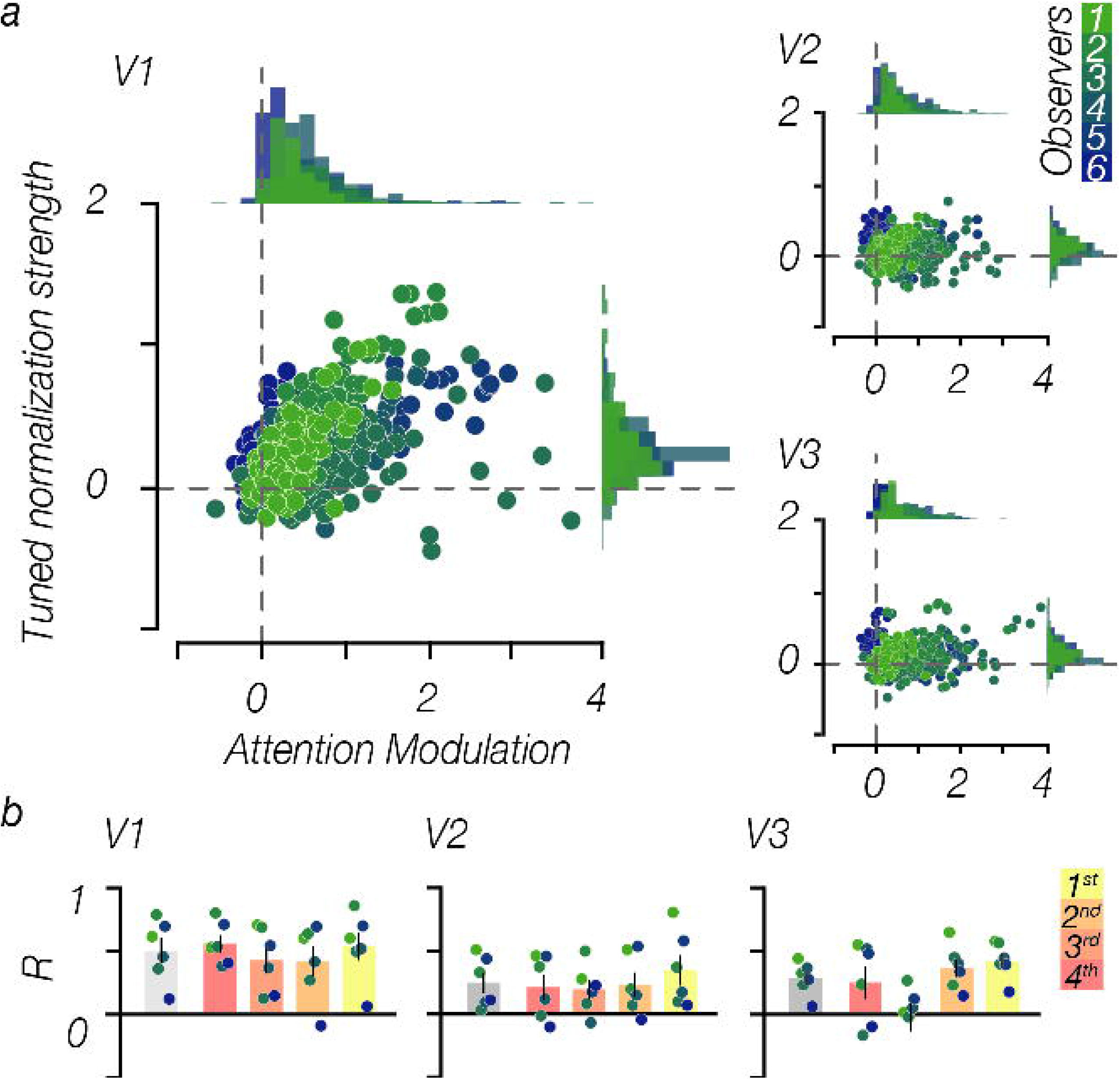
Attentional modulation as a function of tuned normalization strength. a. A tight relationship between tuned normalization strength and attentional modulation is evident for the top 25% selected voxels for each observer. Dots illustrate individual voxels within an area, colors represent an unique participant. b. Spearman correlations were computed for each observer, grey bars represent the mean correlation across observers, while the colored bars represent the correlations when the voxel selection is broken down into 4 bins based on the R^2^ of the independent visual localizer (red: bottom 25%, yellow: top 25% based on the independent localizer scan). Correlations were transformed into a Fisher Z-statistic to allow for statistical comparisons between observers. Error bars denote ± 1 S.E.M.; colored dots illustrate individual observers.

### Tuned normalization modulates spatial attention

To provide converging evidence in support of the underlying relationship between tuned normalization and attention, we carried out an additional experiment, wherein we directly assessed whether attentional modulation is greater when the population response is put under a state of stronger normalization. To do so, we measured BOLD responses for the overlaid stimuli configurations in separate blocks, similar to those constructed in Experiment 1, while covert spatial attention was directed to either the left or right side of fixation. Importantly, to leave enough headroom for an increased BOLD response with attention, we used a lower contrast stimulus (individual components 25% contrast, resulting in a combined overlaid stimulus of 50% contrast). Participants performed a demanding probe detection task, detecting and discriminating whether a probed was embedded in the upper or lower visual field of the attended side of the screen, while maintaining fixation at the center of the screen (**Figure 5a & S3**). This experimental design allowed us to simultaneously measure BOLD responses for either configuration when attention was directed towards or away from the stimulus.

**Figure 5.**
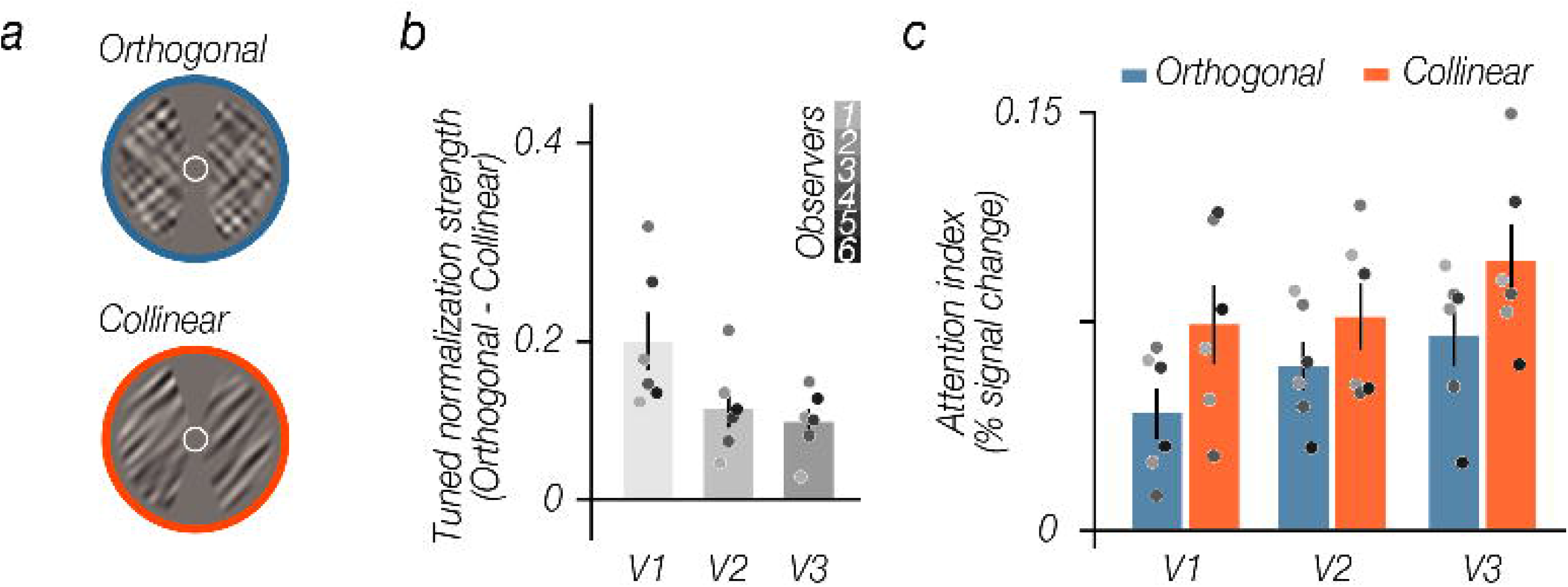
Tuned normalization modulates spatial attention. a. Stimuli were comprised of orientation bandpass-filtered noise gratings (outer diameter 15°; inner diameter 3°; spared midline; individual component was rendered at 25% Michelson contrast, resulting in a combined grating of 50% Michelson contrast), comparable to stimuli used for Experiment 1. Participants performed a demanding spatial attention task, detecting and discriminating a small probe embedded in either the upper or lower visual field of the attended location (a cue presented throughout the block at fixation informed the participant to attend the left or right side of fixation). b. Tuned normalization strength represents the difference between BOLD responses evoked by orthogonal and collinear stimuli blocks when attention was directed away (% signal change). The orthogonal configuration elicited larger BOLD responses compared to the collinear configuration across V1-V3. c. Attention index reflects the difference between BOLD responses when spatial attention was either directed towards or away from the stimuli locations divided by the sum, for both collinear and orthogonal configurations. Error bars denote ± 1 S.E.M., N = 6, grey dots illustrate different participants; stimuli are modified for illustrative purposes.

First, we assessed whether stimuli that share feature information (collinear configuration) yield stronger tuned normalization, compared to stimuli with dissimilar features (orthogonal configuration), when attention was directed to the opposite visual field. In agreement with the results of Experiment 1, we found strong tuned normalization across visual areas (**Figure 5b**; two-sided t-test; V1: *t(5)* = 5.46, *p* = 0.003, V2: *t(5)* = 4.84, *p* = 0.005, and V3: *t(5)* = 5.80, *p* = 0.002). Having established that there is strong tuned normalization when attention is directed away, we then set out to test whether tuned normalization truly dictates the magnitude of attentional modulation. We hypothesized that the largest attentional effects would be evident when a neural population experiences stronger normalization, induced by the similarity between the features of the overlaid stimuli (i.e. attentional effects for collinear > orthogonal).

To quantify the magnitude of attentional modulation, we computed an attention index, which is the ratio of the difference between attending towards vs. attending away divided by the sum of both, for both collinear and orthogonal stimuli configurations. We discovered that attentional effects were indeed the greatest when neural responses were put under a stronger state of normalization (**Figure 5c**; two-sided t-test; V1: *t(5)* = −3.71, *p* = 0.014, V2: *t(5)* = −2.98, *p* = 0.031 and V3: *t(5)* = −2.52, *p* = 0.053). Taken together, these results provide direct evidence to suggest that a more strongly normalized population is more susceptible to an attention-facilitated release from gain control.

### Tuned normalization explains BOLD responses

To parsimoniously describe all the aforementioned findings, we fit our results using an fMRI encoding modeling approach, which incorporates a V1-based energy detection model (**Figure 2b**) to assess the magnitude of tuned normalization (Experiment 1), and whether attention modulates tuned normalization (Experiment 3). We showed earlier that the total contrast energy output of a V1-based energy detection model had the same predicted response energy for both stimuli configurations when standard (untuned) divisive normalization was applied, and that these distributions shifted apart when a tuned component is built into the model, suggesting that tuned normalization could account for the observed differences in BOLD response evoked by collinear and orthogonal stimuli configurations.

Here, we aimed to quantify the contribution of this orientation-tuned component in the normalization model by utilizing a fMRI encoding modeling approach^53,54^ to predict the evoked BOLD responses evoked by each image class. In this modeling approach, we simulated the neural response for a blocked presentation period for both stimuli configurations using a predicted V1-based complex cell output (see **Equation 2**), in response to sets of images within a stimulus presentation block. Note that the model output is believed to be most analogous to striate cortical neural responses^47,48^, and therefore likely most consistent with our V1 data. The time series of the modeled neural response was then transformed into an estimated BOLD response by convolving it with an assumed canonical hemodynamic impulse response function^55^. To estimate the contribution of a tuned component in the normalization pool, we extended the standard normalization model by incorporating this term into the denominator of the model (see **Equation 4**). By optimizing the weights (*ω_or_*) that modulated the contribution of tuned normalization, we were able to obtain simulated block response that best predicted the measured averaged BOLD response to a stimulus block independently for each stimulus configuration (**Figure 6a**). Using this model, the BOLD responses were best explained with a large contribution of tuned normalization for collinear stimuli configurations, while this contribution was much smaller for orthogonal stimuli configurations (two-sided paired t-test V1: *t(5)* = 5.30, *p* = 0.003, V2: *t(5)* = 3.99, *p* = 0.010 and V3: *t(5)* = 3.74, *p* = 0.014; R^2^ model fits collinear configuration: V1 mean = 0.93, SEM = 0.008; V2 mean = 0.94, SEM = 0.005; V3 mean = 0.94, SEM = 0.005; orthogonal configuration: V1 mean = 0.95, SEM = 0.012; V2 mean = 0.96, SEM = 0.007; V3 mean = 0.95, SEM = 0.007; **Figure 6b**).

**Figure 6.**
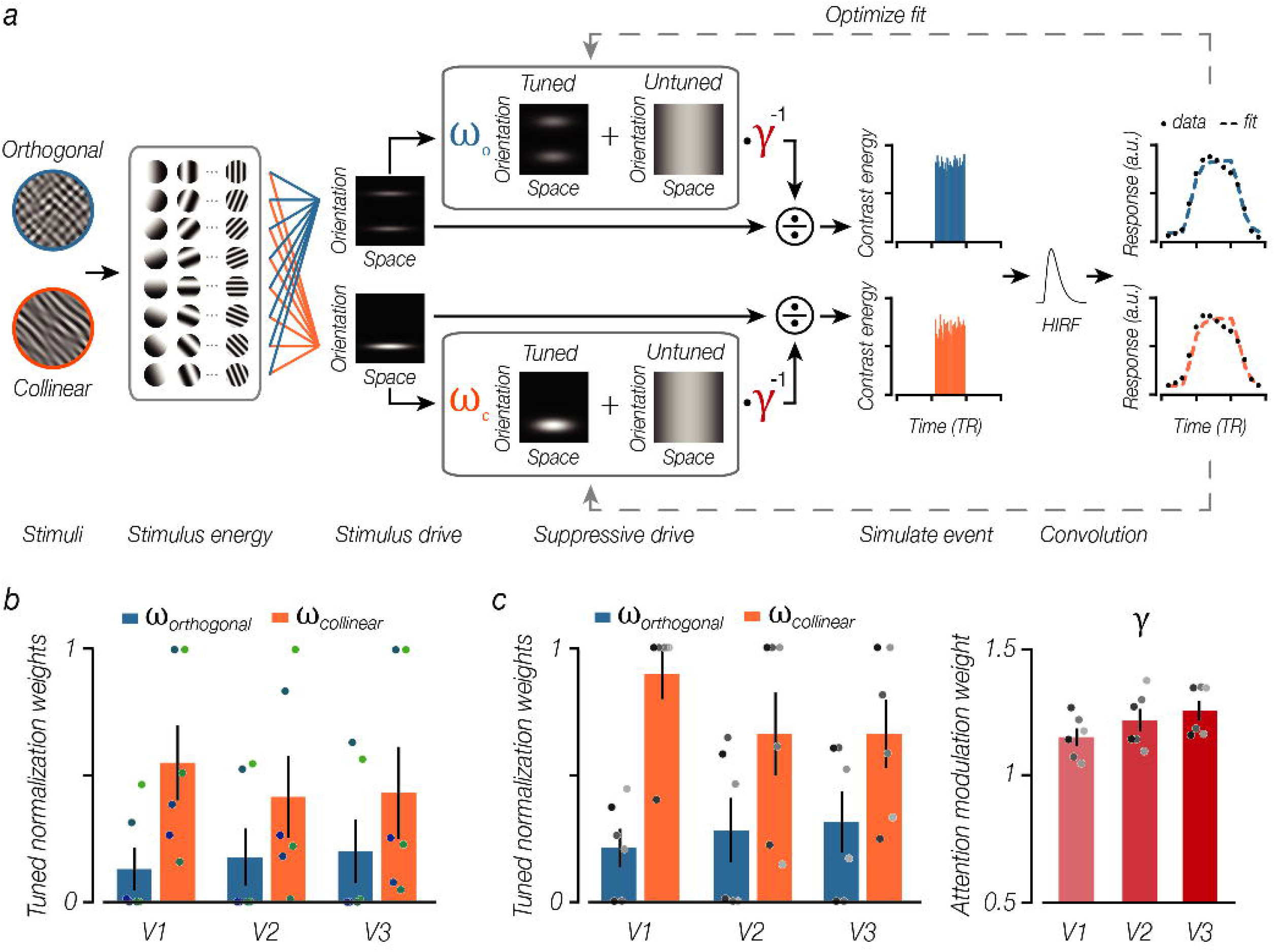
Tuned normalization explains BOLD data. a. fMRI encoding modeling approach utilizes the output of a V1-energy detection model to predict the measured time series to a blocked presentation period evoked by either orthogonal or collinear configurations. The stimulus energy is contained within 8 orientation channels, which undergo divisive normalization to elect a predicted response per presented image. The suppressive pool contains a tuned normalization component, which *ω* modulates (0 = no influence; 1 = full influence), and can be modulated by attention (*γ*> 1, attention results in a release from suppression). b. Estimated weights describing the influence of a tuned component in the normalization model that best predicted the average time series to a blocked presentation period from Experiment 1. c. Left panel: Estimated weights best describing influence of a tuned component in the normalization model when orthogonal or collinear stimuli were unattended. Right panel: Estimated weights describing the magnitude of attention that best predicted the measured BOLD responses in Experiment 3. Error bars denote ± 1 S.E.M., individual dots illustrate different participants.

Armed with a modeling approach capable in summarizing the magnitude of orientation-tuned normalization, we then set out to model the BOLD responses in Experiment 3 – does attentional modulation hinge on tuned normalization? To do so, we built an additional component into the normalization model, allowing for a release from suppression with attention^5^. We simulated the neural response for a blocked presentation period for both stimuli configurations and attention conditions, as described above. We first optimized the weights (*ω_or_*) that modulated tuned normalization based on the unattended BOLD responses, and afterwards we constrained tuned normalization to the best fit, allowing us to predict the influence of attention (*γ*) for each stimulus configuration (**Figure 6a**, see **Equation 5**). Results revealed that the contribution of tuned normalization in the absence of attention for collinear configurations was larger than the contribution for orthogonal configurations (two-sided paired t-test V1: *t(5)* = 8.12, *p* < 0.001, V2: *t(5)* = 5.15, *p* = 0.004 and V3: *t(5)* = 5.07, *p* = 0.004; R^2^ model fits unattended collinear configuration: V1 mean = 0.75, SEM = 0.050; V2 mean = 0.85, SEM = 0.045; V3 mean = 0.85, SEM = 0.044; unattended orthogonal configuration: V1 mean = 0.84, SEM = 0.056; V2 mean = 0.86, SEM = 0.046; V3 mean = 0.85, SEM = 0.046; **Figure 6c**), these estimates are highly comparable to the model estimates obtained from Experiment 1. Importantly, when predicting the average sustained BOLD response when the stimuli were attended, we found that attention modulated tuned normalization via a release from suppression (two-sided t-test different from 1: V1: *t(5)* = 4.46, *p* = 0.007, V2: *t(5)* = 4.94, *p* = 0.004 and V3: *t(5)* = 6.45, *p* = 0.001; R^2^ model fits attended collinear configuration: V1 mean = 0.81, SEM = 0.046; V2 mean = 0.86, SEM = 0.042; V3 mean = 0.86, SEM = 0.040; attended orthogonal configuration: V1 mean = 0.85, SEM = 0.057; V2 mean = 0.85, SEM = 0.041; V3 mean = 0.86, SEM = 0.040; **Figure 6c**).

## Discussion

Taken together, our results reveal that a neural population’s capability for attentional benefits is tightly linked to feature-tuned normalization. The magnitude of attentional modulation depends on the degree to which a population has normalized its response, based on the degree of feature similarity within an image. In the first experiment, we utilized an efficacious method to probe orientation-tuned normalization of population responses within human early visual cortex. By superimposing stimuli that differed in their orientation content, we found that BOLD responses were lower for stimuli that matched in their visual features, compared to stimuli that were comprised of different features. In a second experiment, we found a tight voxel-wise relationship between this measure of tuned normalization strength and an independent measure of attention modulation, suggesting that attention optimizes information processing by modulating divisive normalization. Critically, in a third experiment, we provided direct converging evidence that the magnitude of attentional benefits depends on the degree to which a population has normalized its response; when a neural population was put under a stronger suppressive state, the largest attentional effects emerged. Finally, we propose a tuned normalization model of attention, wherein the incorporation of feature-tuned normalization nicely predicts all of our results, revealing that the degree to which a population exhibits tuned normalization dictates its potential for attentional benefits.

Our results square with a normalization-based model of attention, which posit that attentional modulation arises through interactions with divisive normalization^5,14,31,32^. This model is the prevailing theory, to date, by which attention is believed to act upon neural responses. While previous work provided support for this model^1,28,33,56,57^, our results extend the notion that normalization-driven properties of attention are *feature selective*^10,30,58^. The standard normalization model proposes that the spatial extend of an ‘attention field’ can reshape relative to the stimulus size in order to modulate a population response, and suppression is considered to be feature-agnostic, acting independently from the selective properties computed within a respective region. However, divisive normalization is modulated by contextual influences, where feature similarity results in stronger normalization, a property the model currently does not account for. Previous work has suggested that instead of incorporating a tuned suppressive component into the normalization model of attention, a more parsimonious description to explain differences in the magnitude of suppression is by allowing attention to be feature-selective^26,57^. While the notion of feature-based attention is well established^59,60^, incorporating this into the normalization model does not account for the results we have presented here. In our study, we initially provide evidence that a tuned suppressive component in the normalization model can account for differences between the two stimuli configurations when they were unattended (**Figure 2d**), suggesting that tuned inhibition can arise in the absence of any aid of top-down attentional feedback. Furthermore, in a third experiment we manipulated spatial attention, while holding factors such as stimulus size, attentional window size, and contrast relatively constant, in order to investigate the role that feature-tuned normalization has on attentional modulation (**Figure 6**). Future research measuring the full neural contrast response function, and manipulating features such as the size and shape of the attentional window, will shed more light on precisely how this contribution of feature-tuning is best incorporated into the normalization model of attention.

Normalization is proposed to be a canonical computation throughout cortex and relies on several mechanisms which all serve to regulate the relative strength between neural representations. Two well-established mechanisms within early visual cortex are surround suppression and cross-orientation inhibition^61,62^. Surround suppression is characterized as the modulation of the neural response within the classical receptive field, as a result of the intensity of stimulation presented *outside* of the receptive field^21,63^, while cross-orientation inhibition is the modulation of the neural response induced by presenting two superimposed oriented stimuli components *within* the classical receptive field^62,64,65^. Neuroimaging and psychophysical experiments cannot precisely target a single receptive field, and instead these methods measure population responses evoked by relatively large stimuli, with the spatial area typically spanning far beyond the receptive field of any individual neuron. This likely makes the interactions arising from overlay stimuli, as used here, more analogous to surround suppression. While neuroimaging studies using a typical center–surround stimuli often report an attenuation of the response to the center stimulus^15,38,66^, it is important to consider that this center does not correspond to any particular receptive field center. Instead, the center stimulus drives the response of a large population of neurons, of which only those neurons close to the border between the center and surround stimulus are likely to be attenuated. In this study, we set out to optimize surround suppression by superimposing our stimuli configurations, presented full field (15° visual angle stimulus diameter). We hypothesized that by keeping the orientation of one of the components constant and manipulating the orientation of the second component, we can induce normalization more analogous to surround suppression within all neural populations with receptive fields falling within our stimulus bounds. The superimposed configurations indeed elicited the predicted population responses that one would expect from tuned normalization, as we found lower BOLD responses for those configurations that matched in their orientation content, compared to configurations with orthogonal orientation information. Furthermore, we demonstrated that incorporating a tuned component into the normalization model could account for our results.

Feature-tuned normalization is suggested to play an active role in the efficient coding of natural stimuli^17,20–22,67,68^. Our visual environment is comprised of statistical biases between image features, whereby nearby edges have a higher probability to be co-oriented and belonging to the same contour, as compared to more distant edges^17^. A tuned normalization pool could perhaps incorporate these statistical dependencies by attenuating its strength where features match, leading to an effective boost of the responses at discontinuities, where features are no longer quite as co-aligned. While tuned normalization plays a role by prioritizing processing for salient items, potentially aiding the visual system in segregating figure from ground, we ultimately rely on top-down attentional systems to selectively enhance a small subset of that information for prioritized processing. Selective attention may interact with this figure/ground process by selectively highlighting objects in the environment, which often are defined by their common feature properties.

## Author contributions

I.M.B and S.L conceived and designed the experiments, I.M.B. collected the data. I.M.B conducted data analyses, with assistance from S.L.. I.M.B and S.L. wrote the manuscript.

## Acknowledgements

We thank Louis Vinke & Sara Aghajari for assistance with data collection, Janneke Jehee, Joe McGuire, Rosanne Rademaker and members of the Ling Lab for helpful comments and suggestions. We thank Himanshu Bhat and Thomas Benner (Siemens Healthcare) and Steven Cauley (MGH) for modifying and supplying the simultaneous multislice-BOLD imaging sequence used in this work. This work was supported by NIH EY028163 (SL).

## Methods

### Observers

Six healthy adults participated in the first two experiments (3 male, mean age = 30), and seven adults (2 male, mean age = 28) participated in the third experiment. Five adults participated in all three experiments. All observers provided written informed consent, and had normal or corrected-to-normal vision. The Boston University Institutional Review Board approved the study. One observer who participated in the final experiment was excluded from further data analysis, based on consistent eye-movements towards the cued spatial locations (eye-movement analysis revealed a mean deviation from fixation of >1°). A power analysis indicated that six participants would be sufficient to detect the reported normalization strength and attention effects.

### Apparatus & Stimuli

Stimuli were generated using Matlab (R2013a) in conjunction with the Psychophysics Toolbox^1,2^, rendered on a Macbook Pro (OS X 10.7), and were displayed on a rear-projection screen (subtending ~21°x16°) using a gamma-corrected projector. Participants viewed the display through a front surface mirror. Participants were placed comfortably in the scanner with their heads fixed, using padding to minimize head motion. Stimuli consisted of bandpass-filtered noise gratings (outer diameter: 15°; inner diameter: 3°; at 50% Michelson contrast). The bandpass filter spared only spatial frequencies between 2-3 cycles/degree, orientation content centered at 45° or 135° (orientation bandwidth of 10°), and was smoothed in the Fourier domain to avoid Gibbs ringing artifacts.

### Tuned normalization experiment

Stimuli were the linear combination of the stimuli described above. Superimposing these components created either orthogonal (45°/135° and 135°/45°) or collinear (45°/45° and 135°/135°) stimuli, which resulted in a doubling of the contrast to 100% Michelson contrast.

The two overlaid stimuli configurations were presented at 2Hz (250ms on, 250ms off) for 14s, where each stimulus presentation within a block consisted of unique random noise stimuli. Blocks (2s cue; 14s stimulus presentation) were pseudo-randomized over the course of a run, and interleaved with 16s fixation periods. Throughout the experiment observers performed a demanding fixation task, finding targets in a rapid letter stream presented at fixation (5Hz, letter size: 0.7°). During stimulus presentation blocks, target letters would appear with a probability of 30%, and participants reported whenever they detected a ‘J’ or a ‘K’ amongst distractor letters (**Figure S1**). Observers were capable of discriminating the target letters with high accuracy (mean = 0.93, SE = 0.03).

In addition to the overlaid stimuli, in separate runs we measured the BOLD response to an individual stimulus component (50% Michelson contrast), and doubled this obtained response to create a hypothetical additive sum. Stimuli were presented at 2Hz in blocks (2s cue, 14s stimulus presentation), and were interleaved with baseline periods. Observers performed the same fixation task as described above, reporting the presence of target letters embedded in a rapid letter stream presented at fixation (performance: mean = 0.88, SE = 0.04). Participants completed 5-10 fMRI runs; each run took 272s to complete (8 stimulus blocks per run).

Additionally, a scan session included two visual localizer scans, in which a flickering and rotating contrast pattern was presented within the same aperture as the filtered noise stimuli (blocked presentation, 16s on and off; 6 stimulus blocks per run).

### Attention modulation experiment

To examine the voxel-wise relationship between tuned normalization and attention, we obtained a measure of attentional modulation within the same scan sessions as the previous experiment (n=6). In this experiment, stimuli (single component of the stimuli described above; orientation content centered on 45° or 135°; at 50% Michelson contrast) were presented at 2Hz during a block (2s cue, 14s stimulus presentation). Participants were informed at the start of each stimulus presentation block with a cue (2s) whether to either *attend towards* the stimuli, or to *attend away* from the grating (**Figure S2**). During attended stimulus blocks observers performed an orientation discrimination task, detecting and discriminating a change in the orientation of the stimulus compared to the global orientation (45° or 135°), target stimuli appeared with a probability of 60% throughout the stimulus block. To match task difficulty for the orientation task across observers we titrated individual thresholds to yield an accuracy of 75%. During unattended stimulus blocks observers performed the same fixation task as described above; target letters appeared with a probability of 30%. All stimulus presentation blocks were completely identical, as both orientation and target letters would appear throughout a block, and only the initial cue informed the participant which task to perform (**Figure S2**). The two attentional condition blocks were pseudo-randomized over the course of the run, each interleaved with 16s baseline periods. Behavioral data indicated performance for both tasks was well above chance (attended task: mean = 0.74, SE = 0.05, unattended task: mean = 0.88, SE = 0.04). Participants completed 5-10 fMRI runs; each run took 272s to complete (8 stimulus blocks per run).

### Spatial attention experiment

To directly assess whether attention modulates local gain control, we manipulated covert spatial attention for both stimuli configurations. Participants (n=6) were instructed to maintain their gaze within a fixation circle (diameter, 1°) at the center of the display. Observers viewed stimuli that were the linear combinations of the same bandpass-filtered noise stimuli described above (outer diameter 15°; inner diameter 3°; spared midline), resulting in either orthogonal (45°/135° and 135°/45°) or collinear (45°/45° and 135°/135°) stimuli. Each individual component was rendered at 25% Michelson contrast, resulting in a combined grating of 50% Michelson contrast. Note, the overall contrast of these superimposed stimuli is reduced compared to the previous normalization experiment, as we wanted to leave enough headroom in the BOLD response for the attentional manipulation to take effect.

A cue (2s) at the start of each block informed the participant to allocate their covert spatial attention to either the left or right side of a central fixation point, and remained displayed throughout the block (16s total block duration; **Figure S3**). Observers performed a demanding probe detection task, detecting and discriminating whether the probe appeared at a random location within the upper or lower visual field on the attended side of fixation (probe size 1.5°). Probes could appear on either side of fixation throughout a stimulus block, however observers were instructed to only respond to targets presented on the attended side, as indicated by the cue. Stimulus configuration and attention conditions were counter-balanced and presented in a pseudorandom order, and were interleaved with fixation blocks of equal duration. The behavioral ability to discriminate between targets was comparable for both stimuli configurations, as confirmed by measures acquired outside the scanner (collinear: mean = 0.87, SE = 0.03; orthogonal: mean = 0.90, SE = 0.03; paired t-test: *t(5)* = 0.502, *p* = 0.637). Participants completed 8-14 fMRI runs; each run took 272s to complete (8 stimulus blocks per run). Additionally, a scan session included two visual localizer scans, in which a flickering and rotating contrast pattern was presented within the same aperture as the stimuli (blocked presentation, 16s on and off; 6 stimulus blocks per run).

### fMRI data acquisition and preprocessing

MRI data were acquired at Harvard University’s Center for Brain Science Neuroimaging Center (Cambridge, Massachusetts). Data for the first two experiments were collected in a single scan session, using a 3.0 Tesla Tim Trio MRI Scanner (Siemens, Erlangen, Germany) equipped with a 32-channel head coil. A scan lasted 2h, during which we acquired: an anatomical scan (voxel size: 1.2 mm isotropic) using a T1-weighted multi-echo MPRAGE sequence, and functional volumes with whole brain coverage using a simultaneous multislice (SMS) acquisition protocol (69 slices, TR = 2s, TE = 30ms, flip angle = 80°, FoV = 216mm, voxel size = 2mm isotropic, in-plane acceleration factor 3, multiband factor 3^3,4^. The final experiment was collected using a 3.0 Tesla Prisma MRI Scanner equipped with a 64-channel head coil. A scan lasted 1.5-2h, during which we acquired: an anatomical scan (voxel size: 1.2 mm isotropic) using a T1-weighted multi-echo MPRAGE sequence, and functional volumes with whole brain coverage using a SMS acquisition protocol (72 slices, TR = 2s, TE = 30ms, flip angle = 80°, FoV = 208mm, voxel size = 2mm isotropic, in-plane acceleration factor 3, multiband factor 3^3,5,6^. All analyses were performed in the native space for each participant. Functional volumes were aligned to reconstructed anatomical data, using a surface-based registration between the structural and functional MRI volumes implemented in Freesurfer^7^. Functional data were preprocessed using standard motion-correction procedures, Siemens slice timing correction, and boundary-based registration^7,8^. To optimize voxel-wise analyses, no volumetric spatial smoothing was performed. Robust rigid registration^9^ was performed to align experimental data within each scan session, using the middle time-point of each scan. All further analyses were conducted using custom code written in Matlab.

### Regions of interest

Population receptive field data collected during a separate scan session were analyzed using the ‘analyzePRF’ Matlab toolbox, and used to define regions of interest up to area V3^10,11^. Not all subjects were available for pRF scanning; for one participant in Experiment 1&2 and one participant in Experiment 3, we defined retinotopic regions based on traditional retinotopy scans following standard procedures^12,13^. Within the regions of interest, we defined the top 25% of voxels based on the independent localizer scans (using a standard GLM analysis) for those voxels whose estimated population receptive field (pRF) location fell within the stimulus aperture (15° diameter). This voxel selection ensured that our analysis would be based on voxels that are similarly visually responsive.

### fMRI data analysis

The preprocessed and aligned raw MRI time series per scan, for each voxel, was detrended, high-pass filtered and converted to percent signal change. Task data for all experiments were analyzed by obtaining the activity pattern for each stimulus block, and temporally averaging the BOLD activity across all block of the same condition for every voxel within the ROI, after time shifting by 3 TRs to account of the hemodynamic lag (**Figure S1-3**). For Experiment 1, we quantified the difference between the BOLD response evoked by the orthogonal or collinear stimulus configurations by computing a difference, where a positive difference signals stronger normalization (**Figure 3a**). For Experiment 2, we similarly quantified attentional modulation as the difference between attended vs. unattended blocks (**Figure 3b**).

To compare the degree of voxel-wise dependency between these two measures we computed a Spearman correlation, for all voxels within the defined V1-V3 regions, which was Fisher-Z transformed to allow for comparison between observers (**Figure 4a**). To ensure that the correlations were not driven by a signal-to-noise ratio (SNR) difference, our voxel selection was broken up into 4 equal bins, demonstrating similar correlations within each bin (**Figure 4b**). To ensure that outliers did not drive the computed correlations, we discounted voxels for this voxel-wise correlation analysis that exceeded the mean normalization strength or attentional modulation measure (for each observer) by more than 3 s.d. (1-8 voxels were discounted for each observer within a respective region).

For Experiment 3, covert spatial attention was manipulated to either the left or right side from central fixation, leaving the opposite visual field unattended. This allowed us to examine the effect of attention when attention was either directed towards or away from either visual field, for both collinear and orthogonal stimuli configurations (**Figure S3**). To quantify the magnitude of attentional modulation, we computed a ratio of the difference between attending towards vs. attending away divided by the sum of both (**Figure 5**). This attention index is suggested to be a better representation when comparing an attentional effect between different conditions, as it is not biased by the differences in BOLD responses evoked by the two different stimuli configurations.

### Modeling image statistics

To assess the image statistics of our two stimuli configurations we first analyzed the power of the two image classes in the frequency domain using a standard 2-D Fourier transform. We generated 1000 unique bandpass filtered noise images, which were combined either in a collinear or orthogonal configuration, resulting in 500 overlaid stimuli within each image class (see Apparatus & Stimuli; the size was matched to the screen resolution and visual angle to those images used in the experiments, so that these images were identical to the ones participants viewed in the scanner). Collapsing the Fourier domain power at each frequency band, over all orientations, confirmed that both image types carry the most power (beside the DC component) in those frequency bands the bandpass filter spared (2-3 cycles/°, see above; **Figure 2a**).

Next, we constructed a V1-based energy detection model to describe a plausible underlying neural mechanism that resolves the discrepancy between the image statistics and the evoked BOLD response for each image configuration. We fed the same set of 1000 images used for the Fourier analysis into the V1-based energy detection model consisting of a bank of linear filters^14–19^. Because edge effects can introduce spurious output, we padded each image with 20% of the stimulus size (resulting in an image resolution of 775×775 pixels). The bank of linear filters consisted of 36 spatial frequencies (evenly spaced between 0.5-4 cycles/°), 8 orientations (evenly spaced between 0-180°), 2 phases (0, pi/2), with a receptive field size of 2° visual angle (**Figure 2b**). After convolving each pixel within the image with following filters, we combined the quadrature-phase pairs analogous to a complex-cell energy model^15,20^:

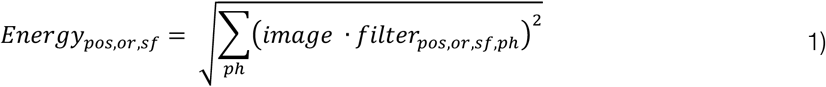

where *Energy_pos,or,sf_* represents the complex cell energy for each image at a given position, orientation and spatial frequency, and *filter_poss,or,sf,ph_* represents the Gabor filter at each particular position, orientation, spatial frequency, and phase.

The contrast energy for all quadrature pairs were summed across spatial frequency scales and averaged over space, resulting in a measure of pooled contrast energy within each orientation channel (**Figure 2b**):

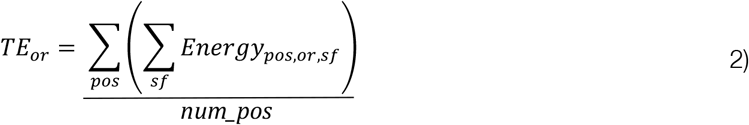

where *TE_or_* represents the total stimulus contrast energy at a given orientation, and *num_pos* reflects the total number of pixels within an image (775×775).

Each images’ summarized complex cell output undergoes untuned divisive normalization, effectively acting as a contrast gain control operator^21–23^:

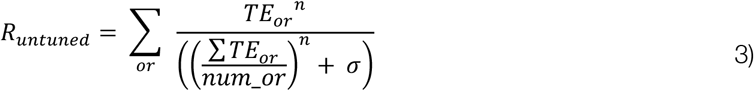

where, *R_untuned_* represents the normalized stimulus energy of each image, *num_or* reflects the total number of orientation channels, ***σ*** is a constant (constrained at 0.5), and ***n*** reflects the nonlinearity in the gain of the response (constrained at 1). For illustrative purposes we computed the total mean stimulus energy over all 1000 images (combining both collinear and orthogonal image configurations) to demean each output, and the maximum stimulus energy over all images to normalize to 1 (**Figure 5c**).

Here, we propose that a normalization pool additionally consists of a tuned component^24^. The untuned component pools equally over all oriented filters (see Equation 3), while the tuned component only contains information in those orientation channels matching the orientation image statistics:

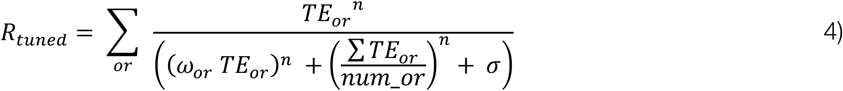

where, *R_tuned_* represents the normalization stimulus energy of each image, *ω_or_* reflects an array of the same size as *TE_or_*, and only those orientation channels matching the stimulus configuration statistics are non-zero, allowing a contribution of tuned normalization (i.e., the tuned component for collinear stimuli could only modulate energy in the 45° orientation channel, while for orthogonal stimuli both 45° & 135° orientation channels were allowed to contribute; **Figure 2d**).

Next, we introduced an fMRI encoding modeling approach^25,26^ to assess the magnitude of tuned normalization that best predicted the BOLD responses for orthogonal and collinear stimuli configurations measured in Experiment 1 (**Figure 6**). We simulated the neural response for a blocked presentation period for both stimuli configurations using the V1-based complex cell output (Equation 2) in response to each image within a stimulus presentation block (**Figure 6a**, 250ms on 250ms off for 14s). The time series of the modeled neural response was then transformed into an estimated BOLD response by convolving it with an assumed canonical hemodynamic impulse response function^27^. To estimate the contribution of a tuned component in the normalization pool, we added this term into the denominator of the model (**Equation 4**). We then optimized the weights (*ω_or_*) that modulated the contribution of tuned normalization, so that the simulated block response best predicted the measured averaged BOLD response to a stimulus block independently for each stimulus configuration, using Matlab’s *fminsearch* function (using nonlinear regression) and custom Matlab procedures (**Figure 6b**).

Armed with a modeling approach capable of summarizing the magnitude of feature-tuned normalization, we then set out to model the obtained BOLD responses in Experiment 3 – does attentional modulation hinge on tuned normalization?^28^ Here, we built an additional component into the normalization model, allowing for a release from suppression with attention:

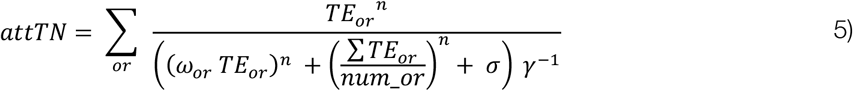

where, *attTN* represents the stimulus energy evoked by collinear or orthogonal stimuli configuration, when attended towards or away from a stimulus. This model is equivalent to Equation 4, but now the normalization pool can be modulated by attention, expressed by *γ*. When *γ* > 1, attention evokes a release from suppression, resulting a larger neural response, and when *γ* = 1 there is no effect of attention. We simulated the neural response for a blocked presentation period for both stimuli configurations and attention conditions, as described above. We first optimized the weights (*ω_or_*) that modulated tuned normalization based on the unattended BOLD responses, afterwards we constrained tuned normalization to the best fit, allowing us to predict the influence of attention (*γ*) for each stimulus configuration.

### Eye-position monitoring

Eye-tracking data were acquired for 5 out of 6 observers for Experiment 1 and 2 (collected within the same scan session), and for 6 out of 7 observers for Experiment 3, using a MR-compatible SR Research EyeLink 1000 system (sampled at 1 kHz). After removing blinks, the mean distance from fixation was computed during time windows corresponding to the stimuli blocks. Specifically, we first calculated the x- and y-deviations, and then for Experiment 1 and 2 computed the absolute distance from fixation, while for Experiment 3 we focused on the x-trace displacement as it gives a better indication whether participants made eye-movements towards the cued attended side. In both scan sessions eye movements were not greater than 0.25° from central fixation (first scan session: 0.24°; second scan session: 0.18°), remaining well within the fixation circle (diameter 1°). However, one observer in Experiment 3 made eye-movements >1° towards the attended side and was excluded from further data analysis. Importantly, for all other observers eye movements did not differ between stimulus configurations, when cued to attend either side of the visual field (repeated measures ANOVA: *F*(1,5) = 0.92, *p* = 0.381).

## Supplementary figures

**Figure S1.**
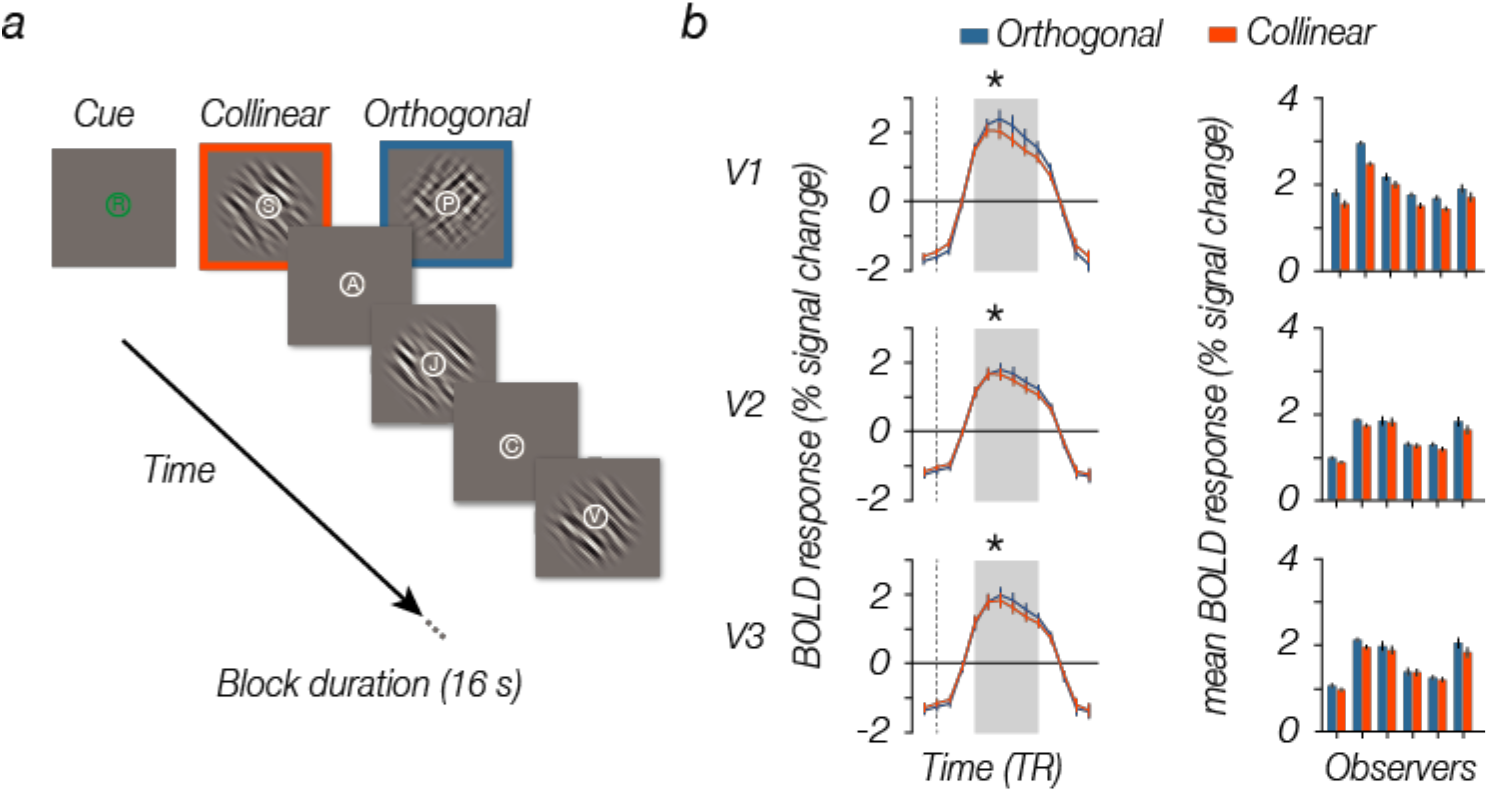
Measuring tuned normalization. **a.** Schematic of an example block sequence. Either collinear (45°/45° or 135°/135°) or orthogonal (45°/135° or 135°/45°) stimuli were presented during a 16 sec block (2s cue period, followed by stimuli presented for 250ms on, 250ms off), while participants performed a fixation task, discriminating target letters in a rapid letter stream. **b.** Mean BOLD responses (left panels) were larger for orthogonal compared to collinear stimuli configurations. Grey highlighted part of the BOLD response reflects the section that contributed to the average BOLD response for each participant (right panels). Stimuli are modified for illustrative purposes; error bars denote ±1 SEM.

**Figure 2.**
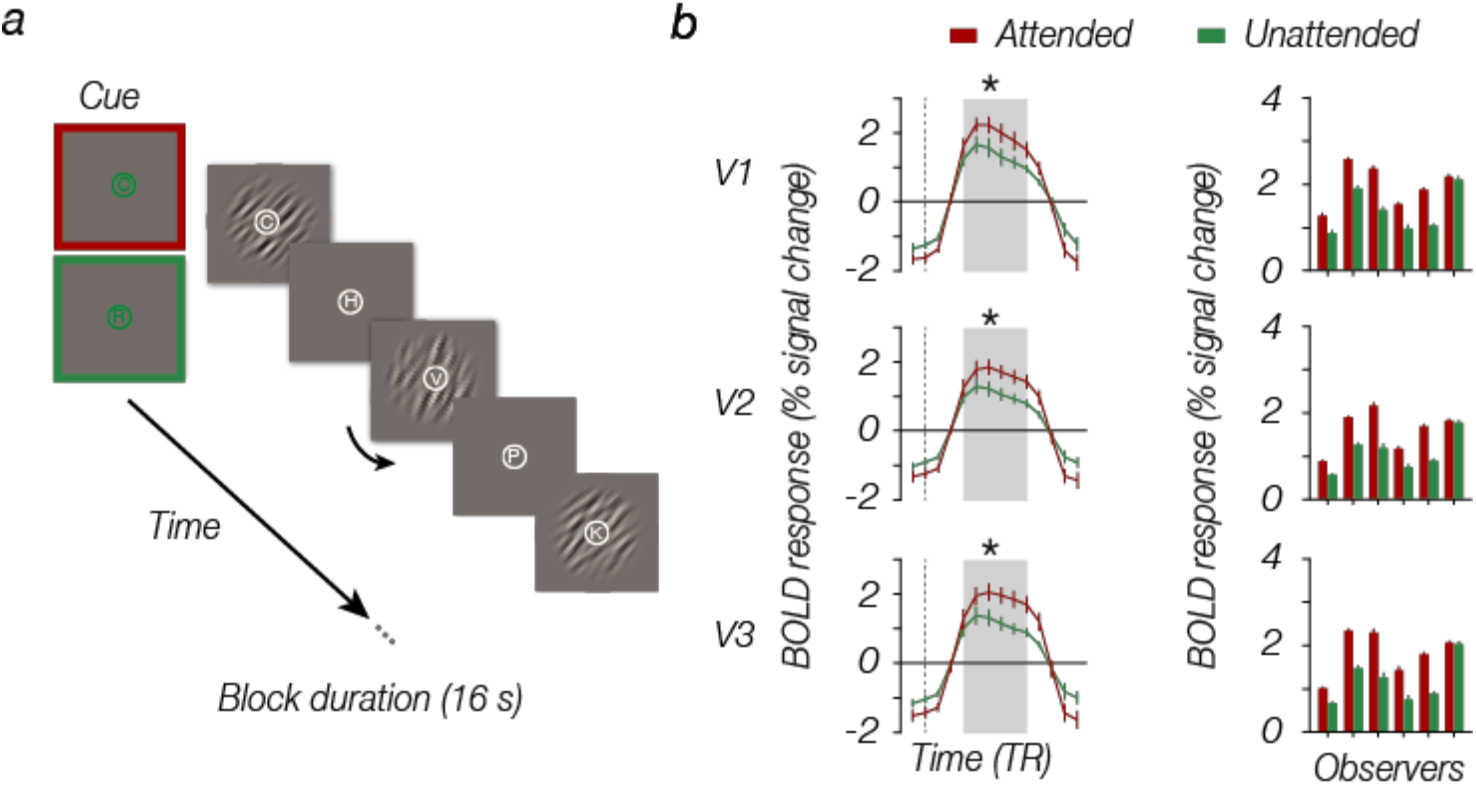
Measuring attention modulation. **a.** Schematic of an example block sequence. A brief cue (2 sec) instructed observers to either attend towards the grating (fine orientation discrimination task), or attend away from the grating (RSVP task at fixation). Both orientation and letter targets would appear throughout a block, only the initial cue informed the participant which task to perform. **b.** Attending towards the stimulus resulted in a larger BOLD response compared to the unattended condition (left panels). Grey highlighted part reflects the section that contributed to the average BOLD response for each participant (right panels). Stimuli are modified for illustrative purposes; error bars denote ±1 S.E.M..

**Figure 3.**
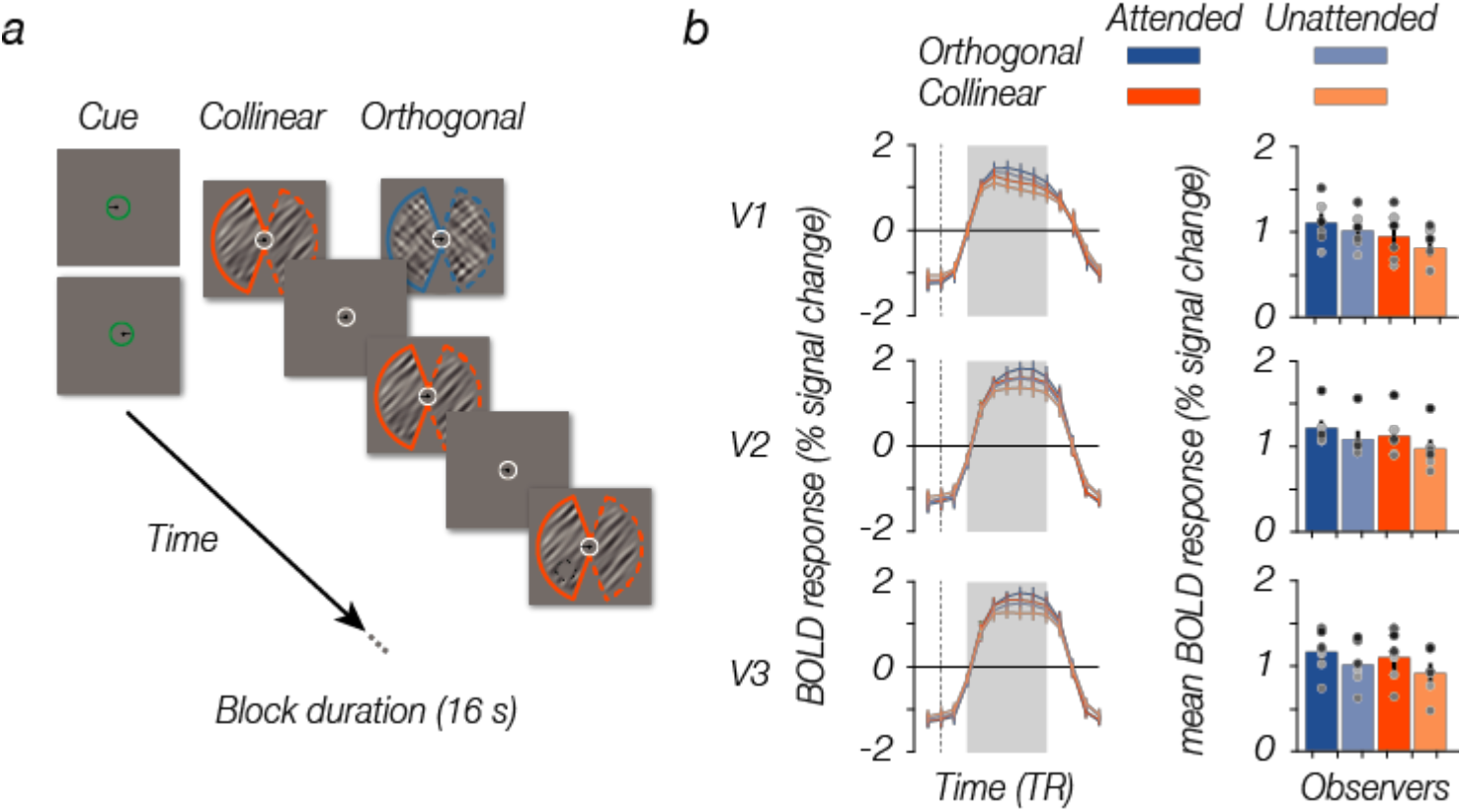
Measuring spatial attention modulation for different states of normalization. **a.** Schematic of an example block sequence. Observers were cued to attend either the left or right side of fixation throughout a block (2 sec), after which either collinear (45°/45° or 135°/135°) or orthogonal (45°/135° or 135°/45°) stimuli were presented (250ms on, 250ms off). Their task was to detect and discriminate whether a target probe (dashed black circle) appeared anywhere within either the lower or upper visual field of the attended stimulus. Dashed orange/blue lines indicate the unattended side, while the solid lines represent the attended side. Attended side and stimulus configuration conditions were counter-balanced and pseudo-randomized throughout a run. **b.** Mean BOLD responses (left panels) for collinear and orthogonal stimulus configurations with and without covert spatial attention. Grey highlighted part of the BOLD response reflects the section that contributed to the average BOLD response for each participant (right panels). Stimuli are modified for illustrative purposes; dots indicate illustrate individual participants; error bars denote ±1 S.E.M..

